# A screen for *Plasmodium falciparum* sporozoite surface protein binding to human hepatocyte surface receptors identifies novel host-pathogen interactions

**DOI:** 10.1101/2020.02.02.929190

**Authors:** Rameswara Reddy Segireddy, Kirsten Dundas, Julia Knoeckel, Francis Galaway, Laura Wood, Gavin J Wright, Alexander D Douglas

**Author notes:** KD and JK contributed equally to this study.

## Abstract

Sporozoite invasion of hepatocytes is a necessary step prior to development of malaria, with similarities, at the cellular level, to merozoite invasion of erythrocytes. In the case of the malaria blood-stage, efforts to identify host-pathogen protein-protein interactions have yielded important insights including vaccine candidates. In the case of sporozoite-hepatocyte invasion, the host-pathogen protein-protein interactions involved are poorly understood. Here, we performed a systematic screen to identify such interactions. We substantially extended previous *Plasmodium falciparum* and human surface protein ectodomain libraries, creating new libraries containing 88 *P. falciparum* sporozoite protein coding sequences and 182 sequences encoding human hepatocyte surface proteins. Having expressed recombinant proteins from these sequences, we used a plate-based assay capable of detecting low affinity interactions between recombinant proteins, modified for enhanced throughput, to screen the proteins for interactions. We were able to test 7540 sporozoite-hepatocyte protein pairs under conditions likely to be sensitive for interaction. We report and characterise an interaction between human fibroblast growth factor receptor 4 (FGFR4) and the *P. falciparum* protein Pf34, and describe an additional interaction between human low-density lipoprotein receptor (LDLR) and the *P. falciparum* protein PIESP15. Strategies to inhibit these interactions may have value in malaria prevention, and the modified interaction screening assay and protein expression libraries we report may be of wider value to the community.

## Introduction

Malaria is a mosquito-borne infectious disease caused by protozoan parasites of the genus *Plasmodium.* Despite some progress in malaria control across the globe, 228 million cases still occurred worldwide in 2018, causing 405 000 deaths, mostly young children in Africa [1]. *Plasmodium* spp. have a complex life cycle that shuttles between the mammalian host and the Anopheles mosquito vector. Malaria infection to the mammalian host is initiated when the mosquito releases sporozoites into the skin while obtaining a blood meal. The deposited sporozoites glide through the skin, enter the bloodstream and traffic to the liver. In the liver, sporozoites actively migrate through several hepatocytes prior to productive invasion, which is characterised by the formation of a specialised compartment, the parasitophorous vacuole (PV). Within the PV, sporozoites undergo several rounds of asexual replication producing thousands of merozoites. Upon rupture of infected hepatocytes, merozoites are released into the blood, invade and replicate inside erythrocytes, leading to the symptoms and complications of malaria [2].

Sporozoite invasion of the hepatocyte is an obligatory step in this life cycle. The *Plasmodium* spp. sporozoite and merozoite share a repertoire of subcellular organelles with each other and with the extracellular stages of other apicomplexan parasites, such as Toxoplasma gondii, reflecting their shared specialisation in the invasion of host cells. A stepwise invasion process appears to be conserved across these parasites, comprising initial low-affinity attachment, calcium-signalling-dependent release of parasite adhesins capable of binding host proteins, formation of a moving junction, and finally actin/myosin-dependent motile invasion into a parasitophorous vacuole [3]. Several parasite ligand – host receptor interactions have been implicated in *P. falciparum* merozoite – human erythrocyte invasion, and blockade of one of these (the interaction of *P. falciparum* reticulocyte binding protein homologue 5 [PfRH5] with basigin) is now a leading vaccine strategy [4]. A number of host and pathogen proteins have been implicated in sporozoite-hepatocyte invasion, as recently reviewed by some of the current authors [5]. Despite this, it is striking that no specific host-pathogen protein-protein interactions have yet been shown to be essential for this process. To our knowledge, no parasite proteins have been shown to interact with hepatocyte proteins such as CD81 and scavenger receptor B1 (SR-BI) [6–9], no host proteins have been shown to interact with parasite proteins such as circumsporozoite protein (CSP), P36 and P52 [9–13], and no clear role in hepatocyte invasion has been shown for the interaction of *P. falciparum* thrombospondin-related anonymous protein (TRAP) with integrin αvβ_3_ [5]. An interaction between host Ephrin type-A receptor 2 (EphA2) and parasite P36 has been suggested but not biochemically demonstrated, and EphA2 has subsequently been shown not to be required for sporozoite invasion [13, 14].

We hypothesized that, like merozoite-erythrocyte invasion, sporozoite-hepatocyte invasion involves multiple protein-protein interactions, identification of which would enable improved vaccine strategies. Biologically important extracellular protein-protein interactions are often of low affinity and can be very transient (for example, the PfRH5-basigin interaction has 0.8 μM affinity and a half-time of 2.7 s). Some of us previously reported a plate-based assay which can be used to identify such interactions, the AVidity-based EXtracellular Interaction Screen (AVEXIS) [15]. We therefore set out to perform an AVEXIS screen to identify sporozoite-hepatocyte protein-protein interactions, modifying the assay method to enhance throughput to a level adequate to investigate a large proportion of the surface protein repertoire of *P. falciparum* sporozoites and human hepatocytes.

## Results

### AVEXIS assay development for systematic high-throughput screen

The AVEXIS assay has previously been used to identify protein-protein interactions, including those with low-affinity and fast dissociation rates [15]. Subsequently we have modified the method to enhance sensitivity by using luciferase rather than β-lactamase as the reporter assay (Galaway *et al*, manuscript in preparation). Here, given the limited prior information about candidate sporozoite ligands and hepatocyte receptors and the large number of expressed candidate proteins, we wished to perform a broad screen and therefore further modified the method to enhance throughput while preserving or improving sensitivity (Figure 1A-B). The assay modifications allowed expression of most proteins in 24-well plates, removed dependency on streptavidin-biotin-mediated capture and so eliminated the need for bait dialysis and expensive streptavidin-coated plates, and retained the sensitive and immediate luciferase-generated luminescence readout.

**Figure 1.**
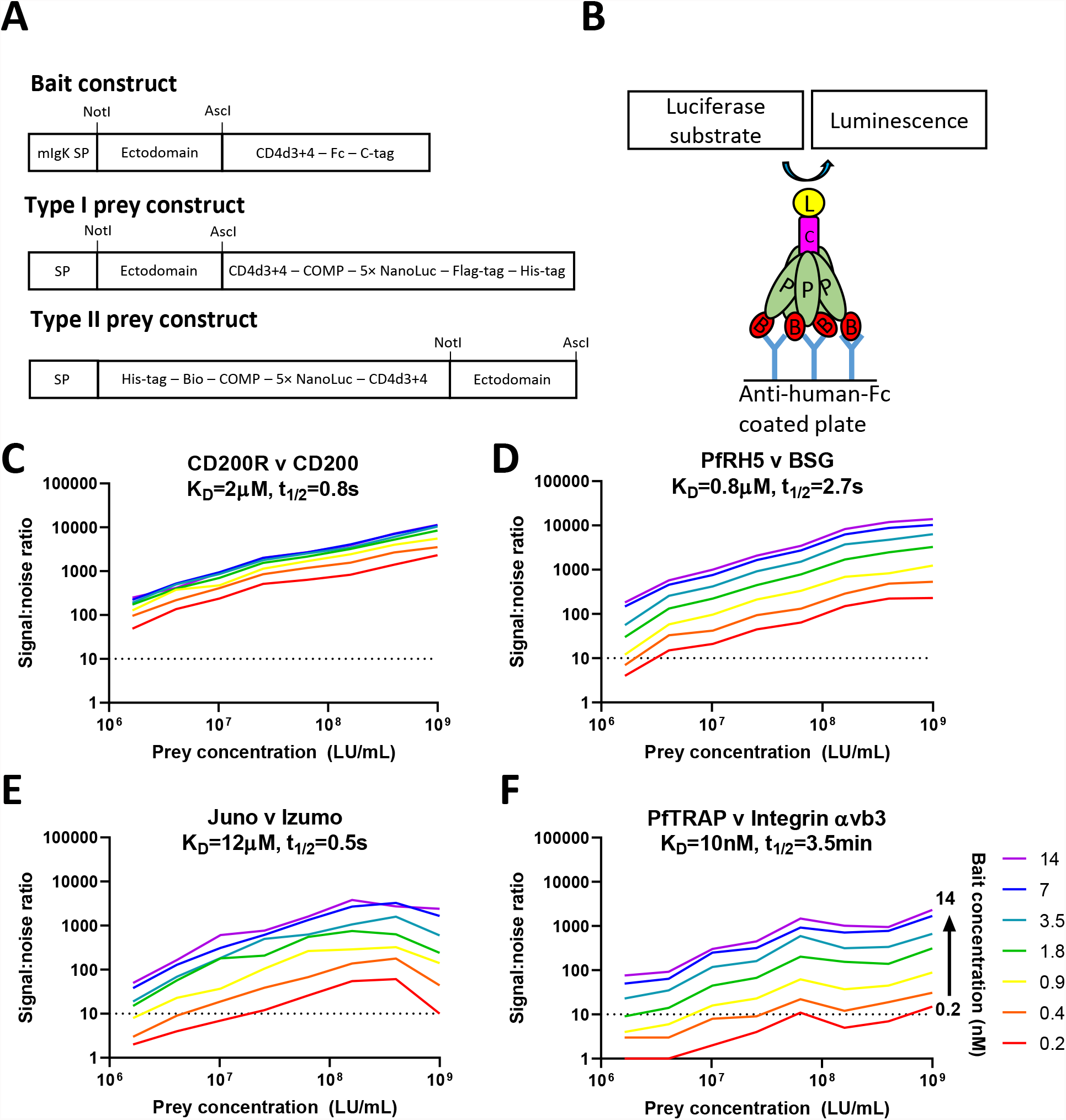
Development and validation of high-throughput modified AVEXIS. **(A-B)** Bait and prey expression constructs and assay schematic. Baits contained a human Fc tag in place of the previously used biotin acceptor peptide. Preys contained a 5×NanoLuciferase tag in place of the previously used beta-lactamase tag. ‘Type I’ and ‘Type II’ preys were used to allow selection of protein orientation (see Methods). Following protein expression, mostly in 24-deep-well plates to enhance throughput, the assay was performed as shown in panel B, with an Fc-tagged bait (labelled ‘B’; red) immobilised on a 96-well plate pre-coated with anti-human IgG-Fc mAb (blue), and probed for interaction with a interacting with a prey (labelled ‘P’; green) protein tagged with pentameric 5×NanoLuc (labelled ‘L’; yellow). The rat cartilaginous oligomeric matrix protein pentamerization domain (labelled ‘C’; pink) mediates pentamerisation. **(C) – (F)** Performance characteristics of the AVEXIS assay were validated using four pairs of proteins with known interaction affinities and dissociation half-lives [5, 16, 17, 48] as indicated on panel labels (for each pair, the bait is named before the prey; for full details, see methods). Graphs depict signal:noise ratio (Y-axis) for each pair at a range of prey concentrations ranging from 1×10^9^ LU/mL to 1.6×10^7^ LU/mL (indicated on X-axis) and a range of bait concentrations ranging from 14nM to 0.2nM (indicated by coloured lines, as labelled on **(F)**). Signal:noise ratios were calculated as described in Methods, by reference to results with the same prey protein/concentration and an irrelevant bait (CD200R, except for CD200 prey, for which Juno was used as irrelevant bait). The dashed horizontal line represents signal:noise ratio of 10.

The modified assay remained capable of the detection of four known protein-protein interactions (Fig 1C-F), including *P. falciparum* RH5: human basigin [16] and mouse Juno: Izumo (which, in monomeric format, is strikingly weak [K_D_ = ~12μM] and transient [t_1/2_ = ~0.5 sec] [17]).

All interactions were detected with a high signal:noise ratio when using bait concentration of 7nM and prey concentration 4×10^8^ LU/mL, and all remained detectable with substantially lower protein concentrations, albeit with lower signal:noise ratios (Fig 1C-F). There was not an obvious relationship between assay sensitivity and interaction affinity or half-life. Sensitivity may instead be influenced by protein quality (e.g. conformational accuracy) and the accessibility of binding sites within these particular constructs.

### Creation of sporozoite and hepatocyte surface protein ectodomain library

Available proteomic, transcriptomic and functional data was reviewed to assemble lists of 84 *P.falciparum* proteins and 189 human proteins which are likely to be expressed on the sporozoite and hepatocyte surfaces respectively, and had architecture amenable to AVEXIS (see Methods). The very large cysteine rich modular proteins CRMP1, 3 and 4 were split into two to three fragments, and three different CSP-based constructs were designed, resulting in a total of 90 sporozoite constructs. The sporozoite protein set partially overlapped with a previously designed library (Knoeckel *et al*, manuscript submitted); 45 of the sporozoite constructs were newly designed for this study. Of the 189 human proteins, 127 constructs were taken from previously designed libraries [16, 18] (and Wright *et al*, manuscript in preparation), while the remaining 62 were newly designed for this study.

Of these constructs, we were able to clone and attempt expression of 88 sporozoite and 182 hepatocyte proteins as AVEXIS baits and preys respectively (i.e. constructs as shown in Figure 1B). Synthesis of the remaining coding sequences was unsuccessful. Full details of constructs produced are provided in Supplementary Tables 1 and 2.

Expression levels obtained and examples of the protein quality as assessed by Western blot are shown in Figure 2 (see also Supplementary Figures 1 and 2, and Supplementary Tables 1 and 2). 82 baits were obtained at concentrations >1 nM which we regarded as potentially informative (based on the results obtained with the test set of known interactions (Figure 1C-F). 61 baits (including CSP, TRAP and P52) were obtained at >7 nM which we regarded as optimal. To achieve these levels, spin filter concentration was required for 54 baits.

**Figure 2.**
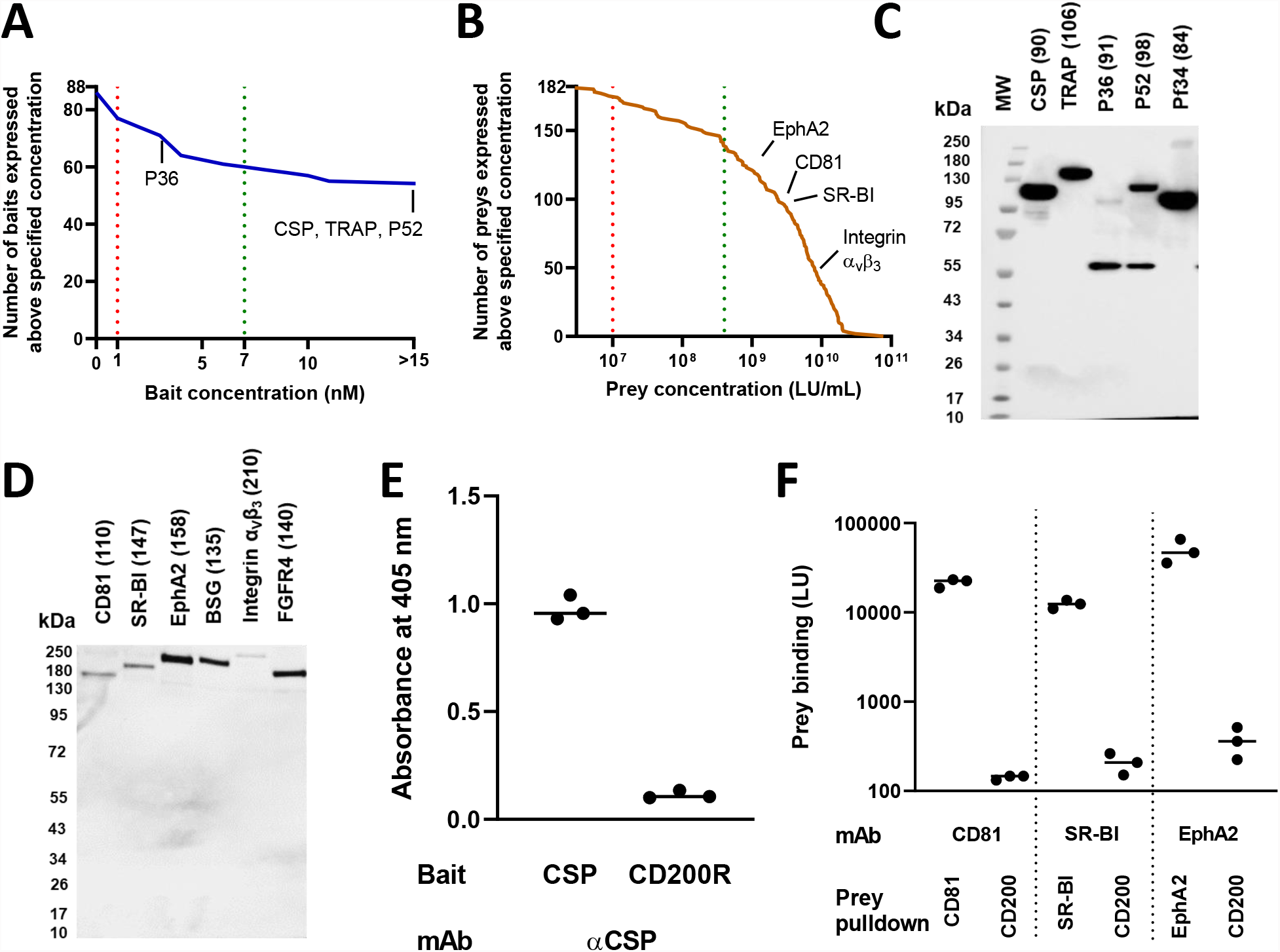
Expression of the P. falciparum sporozoite and human hepatocyte surface protein libraries. Panels **(A)** and **(B)** depict expression levels of bait (by ELISA) and prey (by luciferase assay) respectively, summarised as inverse cumulative distribution functions, and with concentrations of selected proteins of particular interest indicated by name. In the case of 54 relatively weakly-expressed baits and the integrin preys, for which transfections were performed in flasks, the results shown are those obtained after concentration of supernatant. Vertical dashed lines indicate boundaries between optimal, informative and less informative concentrations, as defined based upon the assay validation experiments (see Results and Figure 1C-F). Panels **(C)** and **(D)** show Western blots of selected constructs: CSP, TRAP, P36, P52, Pf34 baits (detected with anti-C-tag antibody) and CD81, SR-BI, EphA2, integrin αvβ3 and FGFR4 preys (detected with anti FLAG-tag antibody; the integrin β-chain is untagged and so not seen). Legend indicates expected molecular weight of each construct in kDa, including tags but excluding post-translational modifications. Some proteins were detected at higher molecular weights than expected, probably due to glycosylation. For Western blots of other proteins, see supplementary figures 1 and 2, and for complete list of baits and preys with details of concentrations and Western blot results, see supplementary tables 1 and 2. The band seen at c. 55 kDa in blots of pre-concentrated baits is believed to represent reactivity of the anti-Ctag antibody with a HEK293-cell protein (rather than degraded bait) as it was also seen in supernatant from cells transfected with irrelevant constructs. **(E)** and **(F)** graphs depict the quantified levels of captured bait and preys using ELISA and luciferase assay respectively.

Of the preys, 165 were obtained at concentrations >1×10^7^ LU/mL which we regarded as potentially informative, of which 139 preys (including CD81, SR-BI, EphA2, and integrin α_v_β_3_) were obtained at >4×10^8^ LU/mL which we regarded as optimal. Results of Western blotting of preys are shown in Figure 2D and Supplementary Figure 2.

To provide additional assurance regarding the quality of key bait and prey proteins, and in particular the activity of the folded proteins in a plate-format assay, we tested whether the full-length CSP bait, and CD81, SR-BI and EphA2 preys could be captured onto 96 well plate using cognate monoclonal antibodies. Captured baits and preys were detected using ELISA and luciferase assay respectively, demonstrating the expected antibody reactivity (Figure 2E-F).

### AVEXIS screen identifies sporozoite ligand: hepatocyte receptor interactions

Having developed the modified high-throughput AVEXIS method and constructed the candidate sporozoite ligand and hepatocyte receptor libraries, we proceeded to a systematic screen for ligand-receptor interactions (Figure 3A and Supplementary Table 3). Signal:noise ratios were calculated by initially correcting for noise attributable to the prey, and then for noise attributable to the bait (see Methods). Raw, prey-corrected and final results are shown on separate worksheets in Supplementary Table 3.

**Figure 3.**
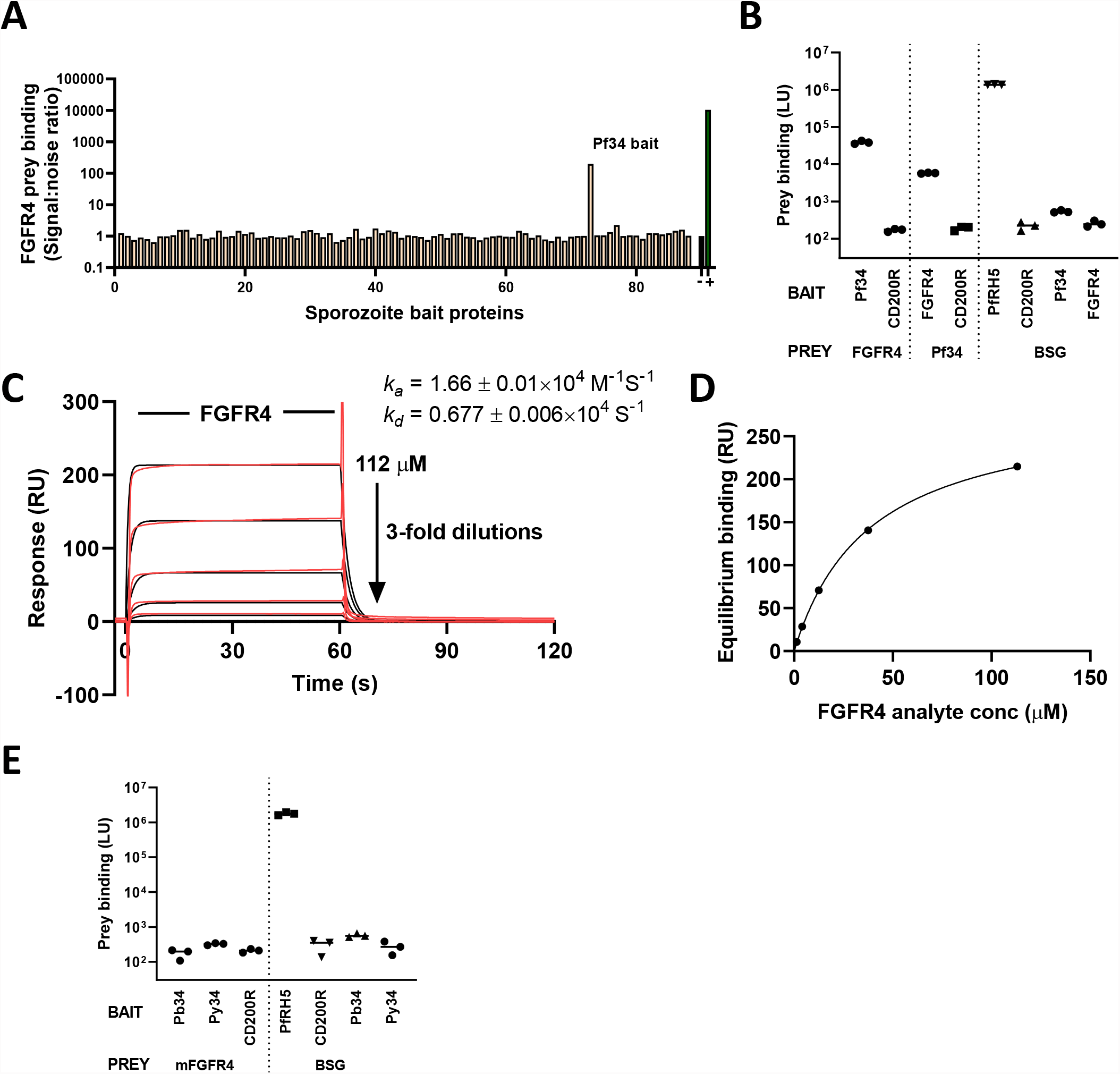
Human FGFR4 interacts specifically with Pf34. **(A)** Human hepatocyte FGFR4 receptor ectodomain prey was tested by AVEXIS for binding to a library of 88 *P. falciparum* ligand ectodomains. Bait numbers correspond to named proteins in Supplementary Table 1. ‘+’ and ‘–‘ indicate PfRH5 bait / BSG prey and CD200R bait / BSG prey, used as positive and negative controls respectively. **(B)** The Pf34-FGFR4 interaction was confirmed by re-testing in three independent experiments, including testing with the reverse orientation of proteins from that used in the screen. Bars represent median ± range of the three experiments. **(C)** and **(D)** show binding of a 3-fold dilution series of FGFR4 analyte (in solution) to Pf34 ligand (immobilised on chip) by surface plasmon resonance. Results shown are representative of two independent replicate experiments with separate protein samples. Panel **(C)** shows kinetics, with observed binding (red lines, double-reference subtracted sensorgrams) overlaid with results of fitting a 1:1 interaction kinetic model (black lines). Panel **(D)** shows equilibrium binding levels (points, from the experiment shown in **(C)**) with the results of fitting a 1:1 equilibrium binding model (line). Curvature indicates binding tending towards saturation, consistent with a specific interaction. **(E)** Absence of AVEXIS-detectable interaction of *P. berghei* and *P. yoelii* orthologs of Pf34 (‘Pb34’ [PBANKA_0721800] and ‘Py34’ [PY17X_0721800], in bait format) with murine FGFR4 (mFGFR4, in prey format).

All 16016 possible sporozoite protein/hepatocyte protein pairs were tested (Supplementary Table 3). Of these, 7540 candidate interactions were tested using protein concentrations we would regard as optimal (bait concentration ≥7 nM, prey concentration 4×10^8^ LU/mL, and good protein quality as assessed by Western blotting), and a further 4718 were tested using protein concentrations which our assay validation data (Fig 1C) suggested would provide a signal:noise ratio of >10 for any of our ‘test set’ of four known interactions.

Across the tested interactions, the highest signal:noise ratio (198) was observed with the combination of the sporozoite protein Pf34 and the human cell-surface protein FGFR4 (Figure 3A). A novel and reproducible interaction was also observed between PIESP15 and LDLR. The protein PIESP15 is under investigation in a separate study by an overlapping team of authors (Knoeckel et al, manuscript in preparation) and so this interaction was not explored further here. Full results of the screen, and a summary table of additional protein pairs with a signal:noise ratio exceeding 5, are presented in Supplementary Tables 3 and 4. No detectable interactions were observed with any of the proteins which have been most prominently implicated as possible sporozoite ligands (CSP, TRAP, P36, P52) or hepatocyte receptors (CD81, SR-BI, EphA2), apart from the known interaction of TRAP with α_v_ integrins.

### Interaction of Pf34 with FGFR4 is specific and consistent with saturable 1:1 binding kinetics

Pf34 (PF3D7_0419700) is a GPI-anchored protein expressed by all parasite stages and localized to rhoptry necks in blood-stage parasites [19]. Pf34 has orthologs in rodent parasites but there are, to our knowledge, no studies to show the role of this protein in sporozoite invasion of hepatocytes. Its *P. berghei* ortholog (PBANKA_0721800) was not covered in recent large-scale ‘PlasmoGEM’ screens of gene essentiality in the *P. berghei* blood, mosquito, and pre-erythrocytic stages [20, 21].

FGFR4 (CD334) is most strongly expressed in the liver, where it is the dominant FGFR family member [22]. The full-length FGFR4 splicing isoform has an extracellular domain which consists of 3 immunoglobulin-like domains, a single transmembrane domain and a cytoplasmic tyrosine kinase domain [23]. A liver-specific signalling pathway through FGFR4, stimulated by ligands including FGF19, is involved in regulation of cholesterol and bile acid metabolism [24]. FGFR4 has previously been implicated in liver stage development of *P. yoelii* in Hepa1-6 cells, as one among a number of ‘hits’ in a screen investigating the role of host kinases in EEF development. [25], although negative results were obtained in a similar screen investigating *P. berghei* [26].

To confirm the Pf34-FGFR4 interaction, the AVEXIS assay was repeated using reciprocally-oriented constructs, with FGFR4 expressed as dimeric bait and probed with pentameric Pf34 prey. Again, clear and reproducible binding was observed (Fig 2B).

To demonstrate Pf34 and FGFR4 interact with the saturable Langmuir kinetics typical of a specific 1:1 interaction, we performed surface plasmon resonance. Weak but clear saturable binding between Pf34 ligand and FGFR4 analyte was observed with an equilibrium binding constant (K_D_) of ~40 μM and rapid kinetics including t_1/2_<1 s (Figure 3C-D, and Supplementary Figure 3). Kinetic values approached the limits for determination using a Biacore T200 instrument and software, and inspection of residual plots suggested that fitted values may have underestimated association and dissociation rates. Re-fitting of the data (using GraphPad Prism) provided a good fit with similar K_D_ (46 μM) and t_1/2_=0.4 s (data not shown). The ability of the AVEXIS assay to detect this extremely weak interaction further illustrates the power of the technique.

Having identified the interaction between human FGFR4 and *P. falciparum* Pf34, we examined whether this interaction is conserved across species by testing murine FGFR4 for interaction with the Pf34 orthologs found in the rodent malaria parasites *P. yoelii* and *P. berghei.* We found no evidence of interaction of either of these protein pairs, despite expression of all proteins at levels in the range expected to give optimal AVEXIS sensitivity (Figure 3E).

Pairwise host-pathogen protein-protein interactions are frequently components of larger multi-molecular complexes, and so we proceeded to investigate possible additional interacting partners of Pf34.

Binding of many FGF family members to their receptors is enhanced by heparin and/or heparan sulfate [27]. Using SPR, we tested whether heparin and heparan sulphate (HS) may have a similar affect upon the Pf34 – FGFR4 interaction. We used a similar design to that used in our experiment measuring Pf34 – FGFR4 kinetics, assessing whether pre-incubation of soluble monomeric FGFR4 with heparin or HS had any effect upon binding to Pf34 immobilised on the chip. Despite using high heparin/HS concentrations (1mg/mL), we did not observe any enhancement of FGFR4 binding (data not shown).

## Discussion

Invasion of hepatocytes by *P. falciparum*. sporozoites is a bottleneck in the malaria parasite lifecycle. Inhibition of this process, particularly by vaccine-induced antibodies, is a major focus in efforts to develop means of malaria prevention. This effort is hindered by limited knowledge of the host-parasite interactions involved in hepatocyte invasion. This study has therefore sought to improve understanding in this area.

Our approach, expressing human and *P. falciparum* proteins in a human cell line and testing them for interaction using a modified AVEXIS assay, was designed to achieve the best sensitivity which we could achieve in a broad, high-throughput screen. Our selection of proteins for study included the majority of single-transmembrane sporozoite and hepatocyte surface proteins for which there is currently convincing proteomic evidence of expression. Transient mammalian cell expression can achieve the same post-translational modifications found in *P. falciparum* proteins. The AVEXIS assay has a strong track record in detection of biologically important interactions, even when they are strikingly weak [15, 16].

Nonetheless, the approach does have limitations. AVEXIS is limited to protein ectodomains and so cannot interrogate many multi-transmembrane proteins (a substantial proportion of the hepatocyte surface proteome). We were unable to express some proteins at all, and a further proportion expressed weakly. Given the high throughput nature of the screen, we cannot be certain about the conformational accuracy of individual proteins, nor the accessibility of potential interaction sites in the constructs. It is likely that relatively weak expression is to some extent an indicator of problematic folding. Thus although results from our ‘test set’ of interactions (Figure 1 C-F) suggest that even very low bait and prey concentrations would in many cases be adequate to detect an interaction, negative results obtained with weakly-expressed proteins are of uncertain reliability.

We found no evidence of interactions with sporozoite proteins of the previously reported host receptors for sporozoite invasion (CD81, SR-BI, EphA2), nor of interactions with host proteins of the suspected sporozoite invasion ligands P36 and P52. The role of these proteins in the invasion process remains incompletely understood.

Our key novel findings are the interactions of Pf34 with the host protein FGFR4 and PIESP15 with LDLR. The interaction of Pf34 with FGFR4 is of very low affinity, but in the context of apposed membranes weak monovalent interactions can sum to provide significant avidity and to become biologically critical (as illustrated by the Juno-Izumo interaction which is essential for mammalian fertilisation [17]). It is also possible that the Pf34 – FGFR4 interaction occurs in the context of a multi-molecular complex which provides higher affinity between the host and parasite members. This possibility is supported by the fact that a co-receptor, β-klotho, contributes substantially to the binding affinity of FGFR4’s endogenous ligands [28]. We were unfortunately unable to express β-klotho in sufficient quantities to further explore this possibility.

The possibility that interactions between Pf34 and FGFR4 could contribute to a functionally-important invasion complex is supported by FGFR4’s role in liver-specific signalling pathways, and evidence of reduced EEF formation in the absence of FGFR4 [24, 25, 29]. Investigation of the effect of disruption of these interactions upon sporozoite invasion is a priority.

## Materials and methods

### Ethics statement

All animal work was conducted in accordance with the U.K. Animals (Scientific Procedures) Act 1986 (ASPA), and the protocols were approved by the University of Oxford Animal Welfare and Ethical Review Body (in its review of the application for the U.K. Home Office Project Licence and P9804B4F1).

### Design of sporozoite and hepatocyte surface protein libraries

Comprehensive lists of sporozoite and hepatocyte surface protein constructs used in this study are provided in Supplementary Tables 1 and 2 respectively.

*P. falciparum* sporozoite surface proteins were selected for study on the basis of their estimated abundance from available surface proteomic and transcriptomic data, plus review of published functional data. In brief, a list of all *P. falciparum* proteins including predicted signal peptides or transmembrane domains was downloaded from PlasmoDB (accessed October 2016). We then used broad criteria to select proteins for manual review. Proteins were considered further if they were among the top 30% of those on the list by abundance in a mass spectrometric analysis of the sporozoite surface proteome [30] or the same authors’ re-analysis of a previous whole-sporozoite proteome [31], or the original report of the sporozoite proteome [32]. Given the poor sensitivity of mass spectrometry for certain proteins, we also included the 5% of proteins on the list for which transcripts were most abundant in two profiles of sporozoite RNA ([33] and RNA-seq data provided by Hoffman *et al* to PlasmoDB.org). We also considered lists of proteins previously found to be present in the micronemes and rhoptries of other *Plasmodium* spp. life-cycle stages [34, 35], and genes for disruption resulted in sporozoite or liver-stage phenotypes, as reported in the RMgmDB transgenic parasite database [36]. These lists were then manually synthesized and reviewed, together with annotation information in PlasmoDB, to compile a set of 79 proteins for which there was reasonable evidence of presence on the surface of the sporozoite (or release to the surface from intracellular organelles during invasion). A further four proteins were added based upon published functional information suggesting a role in sporozoite-hepatocyte attachment and/or invasion (LIMP [37], MB2 [38], LSA-3 [39] and STARP [40]). PfRH5 was included on the basis of its known role in the blood-stage, although we are unaware of any evidence it may have a function in the sporozoite. The total number of proteins selected for study was thus 84. Plasmids encoding 45 of these (though with different tags from our bait configuration) were already available in a previously designed library (Knoeckel et al, manuscript submitted).

For the remaining sporozoite proteins, starting with predicted amino acid sequences from PlasmoDB, SignalP 4.1 and TMHMM v2.0 web servers were used to identify signal peptide and transmembrane domains and hence identify ectodomains. Endogenous signal peptides were replaced with a mammalian signal peptide (from mouse immunoglobulin kappa chain). To avoid inappropriate glycosylation when expressed in mammalian cells, asparagine-X-serine/threonine N-glycosylation sequons were mutated to asparagine-X-alanine. Genes were then codon optimised for mammalian expression. For three of the very large proteins from the cysteine-rich modular protein family (CRMP1, 3 and 4), we designed two to three constructs, together spanning the ectodomain. Given the importance of CSP and its known domain architecture, we designed both a full-length construct and N-terminal and C-terminal domain constructs. The remaining coding sequences were synthesized by Twist Biosciences or, for large or challenging genes, ThermoFisher.

Selection of proteins for inclusion in the human hepatocyte surface protein library proceeded similarly, using available proteomic, transcriptomic and functional data. Our starting point for this was a manually-curated list of human cell-surface proteins. We cross-referenced this with three human primary hepatocyte or hepatoma cell line proteomic data-sets [22, 41, 42] to identify around 1000 cell surface proteins with reasonable proteomic evidence of hepatocyte expression, adding a further 150 proteins which were not detected by mass spectrometry but were abundant in hepatocyte transcriptomes [43]. Because AVEXIS depends upon expression of ectodomains as soluble proteins, we discarded most multi-pass transmembrane proteins from the list, retaining type I, II and III and GPI-anchored single-pass transmembrane proteins, plus a small number of multi-pass proteins with substantial N-terminal ectodomains (typically >100 amino acids). Proteins with extremely large ectodomain coding sequences (>5kb) were also discarded, due to anticipated difficulty of gene synthesis. The resulting list of proteins was manually reviewed and the 189 proteins with the strongest evidence of abundant expression on the hepatocyte surface were selected for study. Plasmids encoding 127 of these (though with different tags from our prey configuration) were already available in previously designed libraries [16, 18] (and Wright *et al*, manuscript submitted). As an exception to our general rule of excluding multi-pass transmembrane proteins, we included extracellular domains of CD81 and SR-BI, as proteins known to have roles in sporozoite-hepatocyte invasion [6, 7, 44]. For CD81, we used the larger of the protein’s two extracellular loops. A similar construct has previously been shown to retain the ability to bind hepatitis C virus E2 [45]. For SR-BI, we used the entire extracellular loop.

For the 62 proteins for which plasmids with coding sequences were not available, hepatocyte protein ectodomain coding sequences were designed as for the sporozoite library, with the exceptions that amino acid sequences were obtained from Uniprot, and that endogenous signal peptides and N-glycosylation sequons were retained. Mouse immunoglobulin κ light chain signal peptides were added to the constructs encoding CD81, SR-BI and type III transmembrane proteins (which lack signal peptides). The library also included constructs encoding a number of integrin heterodimers, as previously described [5].

### Cloning and protein expression

Sporozoite ectodomain coding sequences were cloned using NotI/AscI restriction enzymes into a pTT5-based vector [46], in frame with sequence encoding tags as shown in Fig 1A. Consequently, human Fc tagged bait proteins are expressed as dimers.

Hepatocyte ectodomain coding sequences were cloned similarly into the prey vectors by using NotI/AscI restriction enzymes. Two prey vectors were used, according to the expected orientation of the native protein relative to the cell membrane. The majority of constructs, encoding proteins with free N-termini and with transmembrane domains C-terminal to the ectodomain, were cloned into a ‘Type I’ vector, providing tags as shown in Fig 1A. Type II transmembrane proteins, with free C-termini, were expressed from a ‘Type II’ vector, again as shown in Fig 1A. In the case of integrins, α chain ectodomains were cloned into the Type I prey vector, while β chain ectodomains were expressed without tags.

For recombinant protein expression, ectodomain constructs were transiently transfected using Expifectamine into Expi293F suspension cells as per the manufacturer’s instructions (ThermoFisher). Transient transfections were performed in deep 24 well plates, with 4 mL cells/well and transfected cells were maintained at 37°C, 700 RPM, 8% CO2 and 70% relative humidity. Integrin preys were produced by co-transfection with α and β chain constructs. Supernatants were collected on day 3-4 post-transfection and secreted proteins in the supernatants quantified. Selected baits and preys which were expressed in 4 mL cultures at insufficient levels for AVEXIS were expressed in 25mL cultures in Erlenmeyer flasks, and concentrated by using 30kDa molecular weight cut-off (MWCO) centrifugal filters (Vivaspin, GE Healthcare).

To produce purified monomeric protein for surface plasmon resonance (SPR), a codon-optimised FGFR4 ectodomain coding sequence with C-terminal CD4d3+4-His_6_ tag sequence was cloned into pTT5 [47]. Transfection of Expi293F cells was performed as above. Purification was performed using an AKTA Purifier instrument (GE Healthcare) and HiTRAP Talon column (GE Healthcare). Quality of all purified proteins was confirmed by means of Coomassie staining of an SDS-PAGE gel, demonstrating the expected molecular weight and purity >90% (Supplementary Figure 3).

### Bait and prey quantification and normalisation

Fc-tagged baits were expressed as soluble proteins and quantified by ELISA. For ELISA, Nunc Maxisorp 96 well plates were coated with 50 ng/well of a high affinity mouse anti-human Fc monoclonal antibody (mAb clone R10Z8E9, Abingdon Health) in PBS and incubated overnight at 4°C. Plates were washed 5 times in PBS containing 0.05 % Tween 20 (PBS/T) and once in PBS. Plates were blocked with 5 % skimmed milk in PBS/T for 1h at room temperature. Blocking solution was removed and baits were immobilised onto the plate by incubating for 2h at room temperature. After washing again, 50 μL/well of alkaline phosphatase-conjugated donkey anti-human Fc antibody (Jackson ImmunoResearch, Cat. No. 709-055-098), diluted 1:1000 in PBS was added and incubated for 1h at room temperature. After further washing, 100 μL/well of freshly prepared P-nitrophenyl phosphate substrate (Sigma) in diethanolamine buffer was added and incubated for 10-15 min at room temperature. OD 405 nm was read by a Clariostar plate reader (BMG Labtech) and concentrations of unknown proteins were determined by interpolating from a standard curve of samples of known protein concentration (Fig 2A).

For use in AVEXIS screening, bait concentration was adjusted to a target of 7 nM by dilution with Blocker casein (ThermoFisher) or concentration using a 30kDa MWCO Vivaspin centrifugal filter.

Preys were expressed as soluble 5×NanoLuc tagged proteins. Prey levels were quantified in supernatants using the Nano-Glo Luciferase Assay System according to the manufacturer’s instructions (Promega), with the exception that the substrate solution was diluted 1 in 20 in PBS prior to use. 50 μL of supernatant diluted 1:10,000 with casein was mixed with 50 μL of NanoLuc substrate in a well of 96-well white Maxisorp plate (VWR) and incubated for 3 min at room temperature. Plates were transferred to the Clariostar plate reader and luminescence units (LU) measured (Fig 2B).

All preys were then diluted with 1 volume of Blocker casein (a step which we have found reduces background noise [data not shown]) or, in the case of proteins at ≥8×10^8^ LU/mL, adjusted to a target of 4×10^8^ LU/mL by dilution with Blocker casein. Given that integrin constructs were of particular interest and these preys expressed at relatively low levels, integrin-containing supernatants were concentrated to ≥8×10^8^ LU/mL using a 30kDa MWCO Vivaspin centrifugal filter, prior to addition of casein; other weakly expressed proteins were not pre-concentrated.

### Western blotting

Proteins were mixed with NuPAGE reducing sample buffer and boiled at 70°C for 10 min and separated by SDS-PAGE on NuPAGE 4–12% Bis-Tris gels in anti-oxidant buffer (all from ThermoFisher). Resolved proteins were transferred to a nitrocellulose membrane using the Trans-blot Turbo system (BioRad). Membranes were probed with either biotinylated anti-C-tag conjugate (ThermoFisher, Cat. No. 7103252100) for the detection of baits, or biotinylated anti FLAG antibody (Sigma Aldrich, Cat. No. F9291) for detection of preys. Streptavidin-HRP (Pierce, Cat. No. 21130) was used as a secondary reagent. Proteins on the probed membrane were detected using SuperSignal West Pico Chemiluminescent substrate (ThermoFisher) (Fig 2C-D).

### Antibody reactivity of selected baits and preys

Full length CSP bait was captured from expression supernatant onto a 96 well plate coated with anti-CSP mAb 2A10 (MR4 Resources), with supernatant containing CD200R bait used as a negative control. Captured CSP bait was detected by using ELISA as explained above.

CD81, SR-BI and EphA2 preys were captured from expression supernatant onto a white 96 well plate coated with cognate mAbs (1D6 [Abcam] for CD81, EP1556Y [Abcam] for SR-BI, MAB3035 [R&D Systems] for EphA2), with supernatant containing CD200 prey used as a negative control. Immobilised preys were quantified using luciferase assay as explained above.

### Protein interaction assays by AVEXIS

White Nunc Maxisorp 96 well plates (VWR) were coated with 50 ng/well mouse anti-human Fc monoclonal R10Z8E9 in PBS and incubated overnight at 4°C. Plates were washed 5 times in PBS containing 0.05 % Tween 20 (PBS/T) and once in PBS. Plates were blocked with 5 % skimmed milk in PBS/T for 1h at room temperature. Blocking solution was removed and normalised baits were immobilised onto the plates overnight at 4°C. The next day, plates were washed as above, and normalised preys were added and incubated for 2h at room temperature. Plates were washed and bound preys were detected by using the Nano-Glo Luciferase Assay System (as described for prey quantification).

Given that background levels of signal were observed to vary between preys, the screen was performed by testing one prey against all baits on a single plate. Each plate included the following controls: a ‘pulldown’ of the prey being investigated using anti-FLAG antibody (Sigma Aldrich, Cat. No. F1804), to confirm the addition of functional prey; positive control interaction using *P. falciparum* RH5 as bait and human BSG as prey; and a negative control interaction using irrelevant bait and human BSG as prey.

AVEXIS results are presented in terms of signal:noise ratios, to correct for varying levels of non-specific luminescence attributable to different constructs. The major determinant of the level of noise is the prey (background luminescence after application to wells coated with irrelevant baits varies 10-fold or more between different preys). In the case of experiments using small numbers of baits and preys (<10 in total), signal:noise ratios were thus calculated simply by dividing by the result obtained in wells containing an irrelevant bait and probed with the prey of interest. In the case of the high-throughput screen, we initially performed a ‘single correction’, dividing luminescence by the median result across all baits probed with the prey in question (i.e. the median result on the plate). The use of the median as ‘noise’ is based upon the assumption that the vast majority of protein pairs will *not* interact, and hence the median bait can be regarded as a more representative irrelevant control than any single bait picked to act as a control. A small number of baits appeared to bind non-specifically to multiple preys. We therefore performed ‘double correction’, dividing the ‘single corrected’ result for a given interaction by the median ‘single corrected’ result for all preys tested with the same bait to produce a final signal:noise ratio.

The screen was performed in singlicate. All apparent novel interactions with signal:noise ratios exceeding 30 were repeated, as described in Supplementary Table 4, initially using the same protein preparations. Interactions which were not reproduced were assumed to have been falsely positive in the initial screen, probably due to incomplete plate washing.

Interaction blocking experiments were performed similarly, with the exception that blocking reagents (antibodies) were added onto the immobilised baits on the plate for 1h prior to addition of preys.

### Surface plasmon resonance

Surface plasmon resonance studies were performed at an analysis temperature of 25 °C using a Biacore T200 instrument and HBS-EP+ buffer (both from GE Healthcare). Mouse anti-human Fc monoclonal antibody R10Z8E9 was immobilised onto active and reference flow cells of a Series S sensor CM5 chip using an amine capture kit (both from GE Healthcare). Approximately 1000 response units of Pf34 bait protein was captured onto the active flow cell, with a molar equivalent quantity of PfRH5 bait as a control protein bearing the same tags (i.e. CD4d3+4, human Fc and C-tag).

For use as analyte in the mobile phase, FGFR4 protein bearing CD4d3+4, biotin acceptor peptide and His_6_ tags was purified by affinity chromatography as above. Analytical size exclusion chromatography using a Superdex 200 Increase 10/300 column confirmed the absence of aggregates from the protein preparation (GE Healthcare).

Increasing concentrations of the purified FGFR4 analyte were injected at 30 μL/min over the chip surface, each for 60s followed by a 120s dissociation phase. The Biacore single-cycle kinetics mode (without regeneration between injections) was used, although due to the rapid kinetics of the interaction, all previously bound protein was dissociated prior to the start of each injection.

Data was analysed using Biacore T200 evaluation software (GE Healthcare). All data was double-reference subtracted before model fitting (i.e. both the signal on the reference flow cell with the same analyte and the signal detected during a buffer-only blank injection were subtracted from the signal with the analyte on the active flow cell). 1:1 binding models were fitted to both kinetic data and equilibrium binding data.

Heparin and Heparan sulfate (Sigma) were resuspended in HBS-EP+ buffer and used for surface plasmon resonance studies at 1 mg/mL.

## Acknowledgments

We are grateful to Adrian Hill and Wendy Crocker for support with respect to Optimalvax funding, to Photini Sinnis for advice regarding selection of sporozoite proteins, to Katie Ellis for cell culture assistance, to Simon Draper and David Staunton of the University of Oxford Biochemistry Department Biophysical Instrument Facility for access to Biacore, to Martino Bardelli for provision of the pVIPENTR plasmid, to Adam Ritchie for helpful comments on the manuscript, and to Jake Baum and Joshua Blight for input regarding work with *P. falciparum* sporozoites.

## Supplementary information

### Supplementary Figures

**Supplementary Figure 1.**
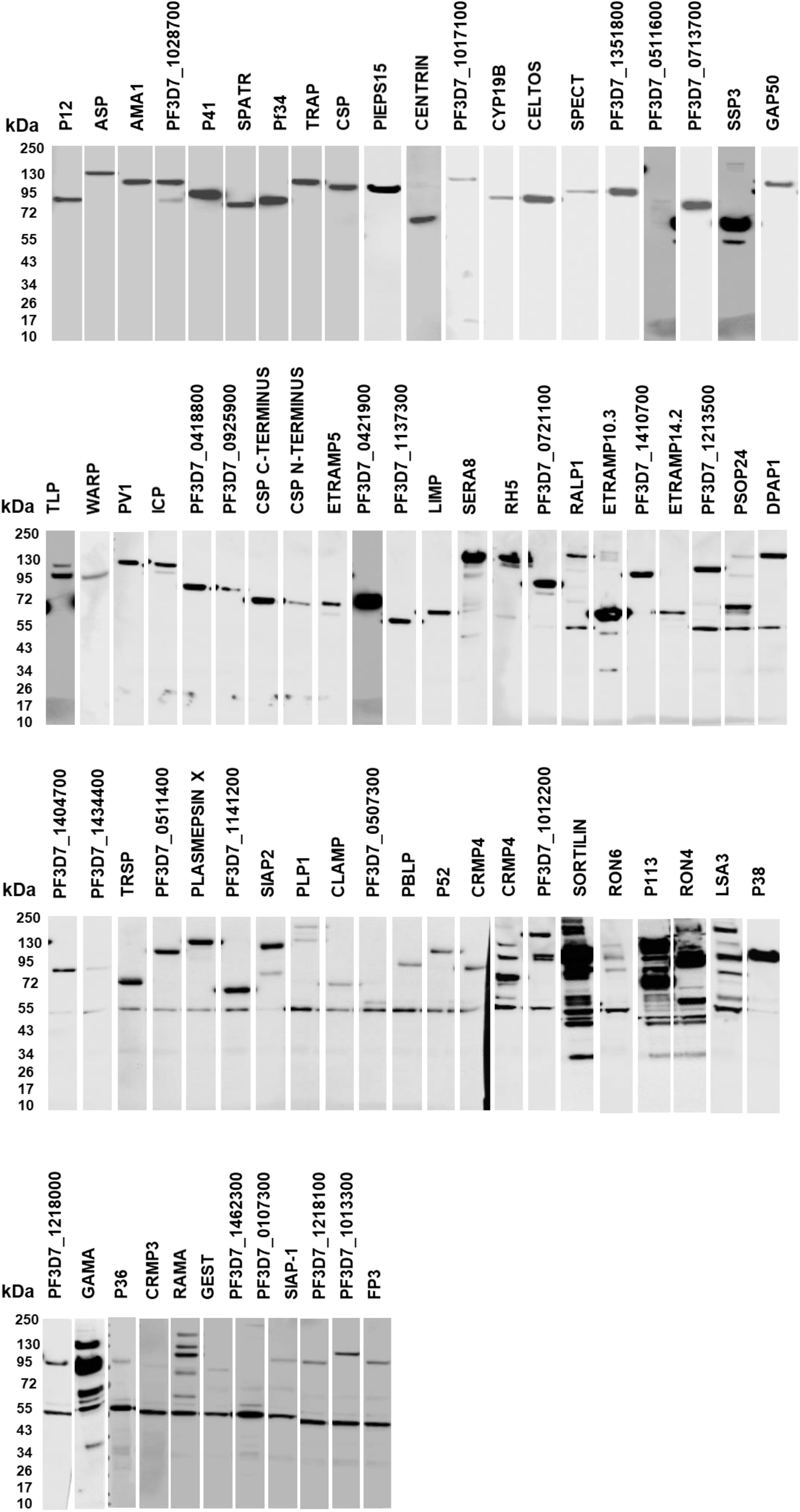
Western blots of sporozoite protein bait library. Proteins are arranged approximately in order of concentration as measured by ELISA, and were detected with anti-Ctag antibody. As described in the text and Supplementary Table 1, expression of 45 of these proteins from related constructs has previously been reported and, in these cases, this figure is intended to demonstrate quality rather than imply novelty. The band seen at c. 55 kDa in blots of pre-concentrated baits is believed to represent reactivity of the anti-Ctag antibody with a HEK293-cell protein (rather than degraded bait) as it was also seen in supernatant from cells transfected with irrelevant constructs.

**Supplementary Figure 2.**
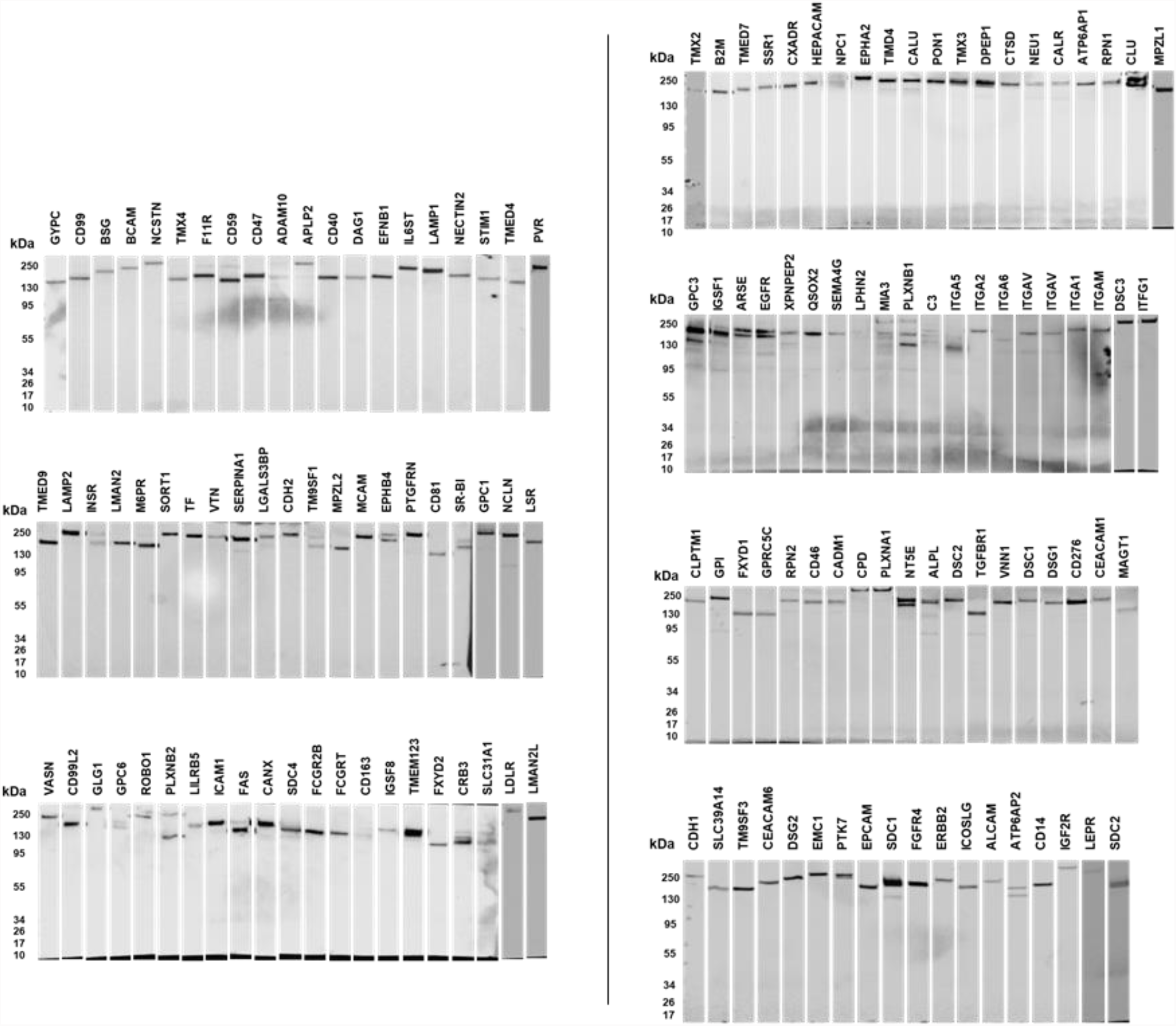
Western blots of hepatocyte surface protein prey library. Proteins are arranged approximately in order of concentration as quantified by luminescence measurement, and were detected with anti-FLAG-tag antibody. As described in the text and Supplementary Table 1, expression of 127 of these proteins from related constructs has previously been reported and, in these cases, this figure is intended to demonstrate quality rather than imply novelty.

**Supplementary Figure 3.**
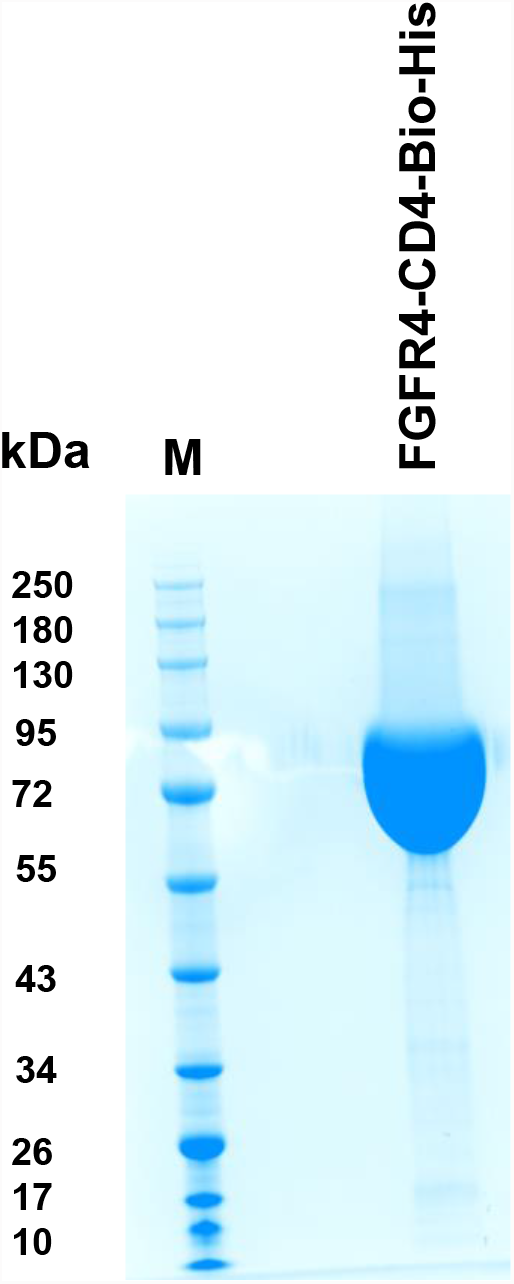
Quality of purified protein used for SPR. Purified monomeric FGFR4-CD4-Bio-His protein (~65 kDa) used for SPR, on SDS-PAGE gel stained with Coomassie Blue.

### Supplementary Tables

**Supplementary Table S1. *S*porozoite protein ectodomain library details**

Details include bait index number (corresponding to numbering in Figure 3A), construct boundaries and sequence, and expression levels (corresponding to Figure 2A).

**Supplementary Table S2. Human hepatocyte protein ectodomain library details.**

Details include construct boundaries and sequence, and expression levels (corresponding to Figure 2B). Integrin α and β chain constructs are listed on the second worksheet, and the results of integrin heterodimer expression are listed on the third worksheet.

**Supplementary Table S3.**

First worksheet presents complete AVEXIS screen results, presented in terms of double-corrected signal:noise ratio (see Methods). Color scale denotes the signal:noise ratio of each interaction, ranging from dark green (low) through yellow to dark red (high).

Second and third worksheets show results with FGFR4 prey against all sporozoite baits (as shown in Figure 3A) and with Pf34 prey against all sporozoite baits (as shown in Figure 4B).

**Supplementary Table S4.**

All protein pairs with a signal:noise ratio exceeding 5 in the initial AVEXIS screen.

